# Methods for analysing lineage tracing datasets

**DOI:** 10.1101/2020.01.11.901819

**Authors:** Vasiliki Kostiou, Huairen Zhang, Michael WJ Hall, Philip H Jones, Benjamin A Hall

## Abstract

A single population of stem cells maintains many epithelial tissues. Transgenic mouse cell tracking has frequently been used to study the growth dynamics of competing clones in these tissues. A mathematical model (the ‘single progenitor model’) has been argued to reproduce the observed stem cell dynamics accurately. This requires three parameters to describe the growth dynamics observed in transgenic mouse cell tracking- a division rate, a stratification rate, and the probability of dividing symmetrically. Deriving these parameters is time intensive and complex process. We compare the alternative strategies for analysing this source of experimental data, identifying an approximate Bayesian computation-based approach as the best in terms of efficiency and appropriate error estimation. We support our findings by explicitly modelling biological variation and consider the impact of different sampling regimes. All tested solutions are made available to allow new datasets to be analysed following our workflows. Based on our findings we make recommendations for future experimental design.

## Introduction

The stem cell dynamics of squamous epithelial cells is a major subject of study in biomedicine. Squamous epithelial tissues cover the external surface of the body, the mouth, and the oesophagus. Importantly, most common human cancers develop from these tissues. Understanding the rules of cell fate decision is therefore fundamental to explain not only healthy tissue growth and maintenance but also the mechanisms of wound healing, mutagenesis and cancer. Epithelial tissues consist of layers of keratinocytes, and in mice, skin and oesophageal tissues are maintained by a single layer of cells at the base of the tissue. Progenitor cells in this basal layer stochastically differentiate, cease cell division, and then stratify into the upper layers of the tissue, migrating to the surface before eventually being shed (Fig. 1a). Human squamous tissues have a more complex organization but share several features and are believed to be maintained in a similar manner.

**Fig. 1:**
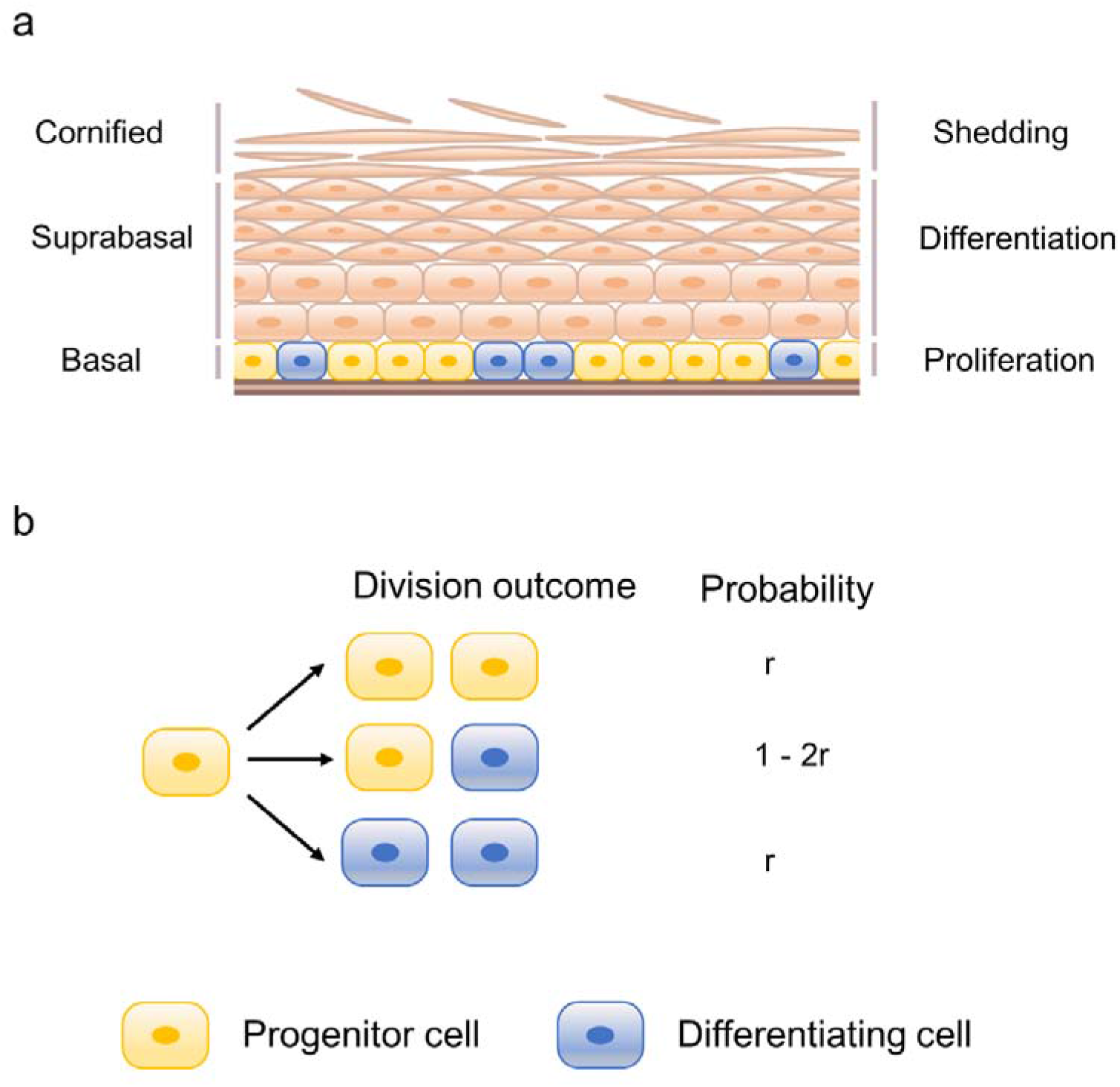
The architecture and maintenance of murine stratified squamous epithelial tissues. a) Proliferation is restricted to the deepest basal layer. Upon differentiation, basal cells exit the cell cycle and migrate through suprabasal layers, until eventually, they reach the surface where they are shed from the tissue. Cell production and loss should be perfectly balanced so that homeostasis and tissue proper function is achieved. b) According to the single progenitor model, stratified epithelial tissues are maintained by a single, equipotent population of progenitor cells which divide stochastically to generate either two proliferating daughters, two differentiating daughters with equal probabilities or one daughter of each type.

The ‘single progenitor’ model (Fig. 1b) has been shown over several studies to accurately describe the observed progenitor cell dynamics in transgenic lineage tracing experiments (1–4). Using three parameters, the division rate, the stratification rate, and the probability of symmetric division, this model predicts average clone size, clone size distributions, tissue homeostasis, and clone survival probabilities. Whilst simulation based techniques have been used to fit experimental data, an analytical solution has also been described (5) allowing for maximum likelihood calculations. Model parameterization and accurate representation of uncertainty however remains problematic, particularly in light of short-term data from in vivo histone dilution assays and live imaging that show cell cycle distribution times do not follow the exponential distributions assumed in the analytical model (6–7).

Here we report the results of exploring alternative approaches applied to the analysis of both published and synthetic datasets. We find that simulation-based maximum likelihood methods require extensive sampling to find a distribution of parameters making them intractable for many analyses. We further find that the use of the published analytical solution allows identification of a single parameter with narrow confidence intervals. However, the analysis of synthetic datasets with realistic cell-cycle distribution times and biological variation between samples suggest that these intervals are too narrow and do not accurately reflect uncertainties in the method and underlying data. We go on to show that an Approximate Bayesian Computation (ABC) based approach using a non-Markovian simulator gives appropriate error bars at an acceptable computational cost. All code underlying this is made available as a Python notebook, enabling the easy analysis of newly collected datasets. Finally, we use synthetic data to explore the relationship between parameterization and the methods of data collection, concluding that single timepoints with typical sampling from the literature (three mice) show high variability that when analysed in isolation could be open to misinterpretation.

## Results

### Approaches based on approximate Bayesian computation more accurately estimate parameters and uncertainties than maximum likelihood

In order to identify the most appropriate strategy for parameterising progenitor cell clonal dynamics we explored the effectiveness of different inference techniques. Maximum likelihood approaches have been widely used in previous publications (3,4,8). In this approach to the analysis of transgenic lineage tracing, the likelihood of different parameter combinations is estimated from the frequency at which different clone sizes have been observed at different timepoints, and a calculation of the probability of a given clone size arising. This probability can be calculated using a published analytical solution of the branching process describing the single progenitor model (5). This however assumes that cell cycle times are exponentially distributed, which is not the case and can undermine this analysis (6). In this situation, bespoke simulation tools need to be run with extremely high sampling (over one-hundred thousand simulations per parameter combination) in order to calculate the clone size probability distributions accurately, at substantial computational cost. The estimates of likelihood calculated from the observed clone sizes and probability distributions can then be used to calculate the most likely combination of parameters and confidence limits.

An alternative approach, Approximate Bayesian Computation, does not rely upon the calculation of the probability of clone sizes. Instead, models with specific parameter sets are simulated and the resultant distributions compared with experimental observations. As this approach is simulation based, it is insensitive to the use of non-Markovian processes. Using a distance metric such as the inter-quantile distances between distributions, or the Kolmogorov–Smirnov statistic, sets of parameters can be collected that have similar properties to the experimental observations. Sequential Monte Carlo Approximate Bayesian Computation (SMC-ABC) is a specific ABC protocol that can rapidly calculate acceptable sets of parameters through iteratively identifying an “acceptable” set of parameters (those below a specific distance), and then perturbing the parameters, simulating models based on those parameters, and rejecting those that fail to meet a new lower distance threshold until a new set of accepted parameters is defined. Through repeated rounds of testing, perturbation, and rejection populations of parameters can be identified that fit the data increasingly well, and can be used to identify both uncertainty and best fitting parameters (9–10). This approach is continued until the rejection rate starts to rise, at which point it is believed that overfitting starts to occur (9).

The above three methods (MLE with analytical solution, MLE with simulation, and ABC) were tested on the analysis of both experimental (3) and synthetic datasets with exponentially distributed cell cycle times. These simulated datasets were generated based on the clonal data provided in (3), and allowed us to use an increased sampling (100,000 total simulated clones) compared to the typical sampling followed in the experimental protocols (three animals per timepoint, one hundred clones per animal). All approaches broadly agreed on the estimated parameters for both experimental (Fig. 2a-c) and synthetic data (Fig. 2d-f). MLE using the analytical solution was able to infer the expected parameter values efficiently (*r* = 0.064, *ρ* = 0.68), producing narrow confidence intervals (*r* = (0.05, 0.068) 95% *CI*, *ρ* = (0.64, 0.72) 95% *CI*) and a smooth likelihood distribution when applied to the experimental dataset (Fig. 2a). In contrast, MLE using model simulations suggested that a sample size larger than 100,000 clone simulations per parameter set would be required to produce an appropriate distribution, increasing computational effort substantially. Simulations of 100,000 clones for 19×19 parameter combinations estimated similar parameter values (*r* = 0.06, *ρ* = 0.64), but produced unrealistically narrow confidence intervals of zero width and the shape of the likelihood distribution was no longer smooth suggesting that sampling undermined the analysis (Fig. 2b). The increased computational demand required for this analysis (28 hours CPU time for 19×19 combinations compared to 1.3 hour using the analytical solution for the same grid), restricted the number of parameter combinations that were searched.

**Fig. 2:**
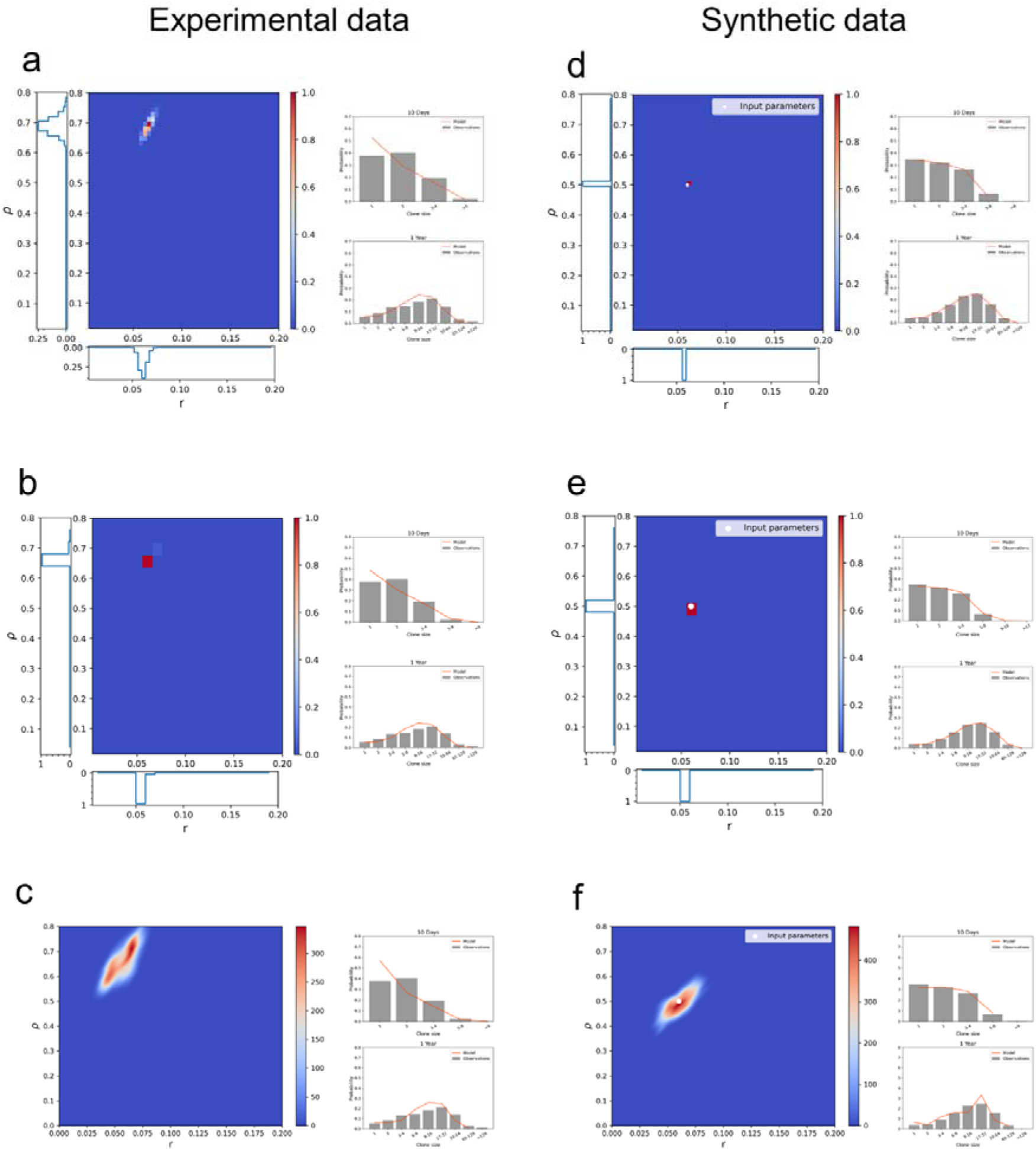
Analytical solution and simulation-based methods inferring single progenitor parameters from both lineage tracing and synthetic datasets. The three different parameter estimation strategies applied to mouse oesophagus lineage tracing data from (3) (a-c) and synthetic datasets generated by performing single progenitor model simulations with parameters λ=2.9/week, r=0.06, ρ=0.5 and assuming exponentially distributed cell cycle time (d-f). a, d) A maximum likelihood approach based on the analytical approach gives a narrow distribution of likelihoods for each parameter. b, e) With 100,000 simulations per parameter set, inferred likelihoods are noisy and fail to give smooth distributions in r or ρ. c, f) SMC-ABC running for 10 generations. Heatmap plots Kernel Density Estimate of final population of parameter sets. a, b, d, e: Heatmap shows the likelihood distribution, whilst likelihood estimates for each parameter are shown alongside. a-f Right: Basal clone size probability distributions of the input data (grey) and the inferred parameters (orange) as obtained at an early and late timepoint.

In contrast to both MLE approaches, the SMC-ABC approach produced a smooth distribution (Fig. 2c) with substantially larger confidence intervals when used to analyse the experimental data (*r* = 0.06 (0.04, 0.073) 95% *CI*, *ρ* = 0.71 (0.56, 0.77) 95% *CI*). Whilst the peaks of the MLE analysis were within these intervals, the distribution of acceptable parameters was offset relative to the MLE likelihoods. Given that it is known that there is substantial biological variation (divisions times are estimated to vary by around 10 % (6)), this raises the question of whether MLE approaches were underestimating the uncertainty that arises due to biological variation. Alternatively, the analysis of synthetic data, which is based on a single parameter, and hence has no uncertainty from biological variation (Fig. 2d,e,f), may suggest that uncertainty is overestimated in ABC, where broad CI were observed. Furthermore, a known issue of the analytical solution is the assumption of an exponential distribution of stem cell cycle times. Both histone dilution and live imaging experiments on epithelial tissues have shown that there is a refractory period in which no division can occur (6–7). This can undermine the analysis leading to the identification of incorrect parameters (6). The issue in assuming an underlying exponentially distributed cell cycle time is also highlighted in the obtained clone size distributions at early time points, where the difference to the clone size distributions of the experimental data is higher compared to later time points (Fig. 2a-c, Supplementary Figures. 1,2).

To test the influence of both cell cycle distribution times, and biological variation, both features were included in models used to generate a new set of synthetic data. These new synthetic datasets were generated with Gamma distributed cell cycle times (Gamma distribution parameters taken from (6)). Furthermore, *r*, *ρ* and *λ* parameter values were drawn from a normal distribution around *λ* = 2.9/week ± 0.1 *SD*, *r* = 0.09 ± 0.01 *SD*, and *ρ* = 0.7 ± 0.05 in order to introduce noise and mimic the variation observed in real datasets. Where the analysis allowed, we took account of the shape of the gamma distribution. This can be inferred through an orthogonal experimental protocol and so was not fit alongside other parameters (6).

Application of the analytical solution to the realistic synthetic datasets revealed that the estimated likelihoods were both inaccurate and overly precise (Fig. 3a). The introduction of more non-Markovian cell cycle times leads to misleading calculated parameters in the analytical MLE approach (Fig. 3a). Moreover, the narrowness of the confidence intervals calculated with this method excludes the true parameters. We suggest that this underestimation of uncertainty arises as a result of limited sampling, and this same issue has been noted in other, unrelated systems (11–12).

**Fig. 3:**
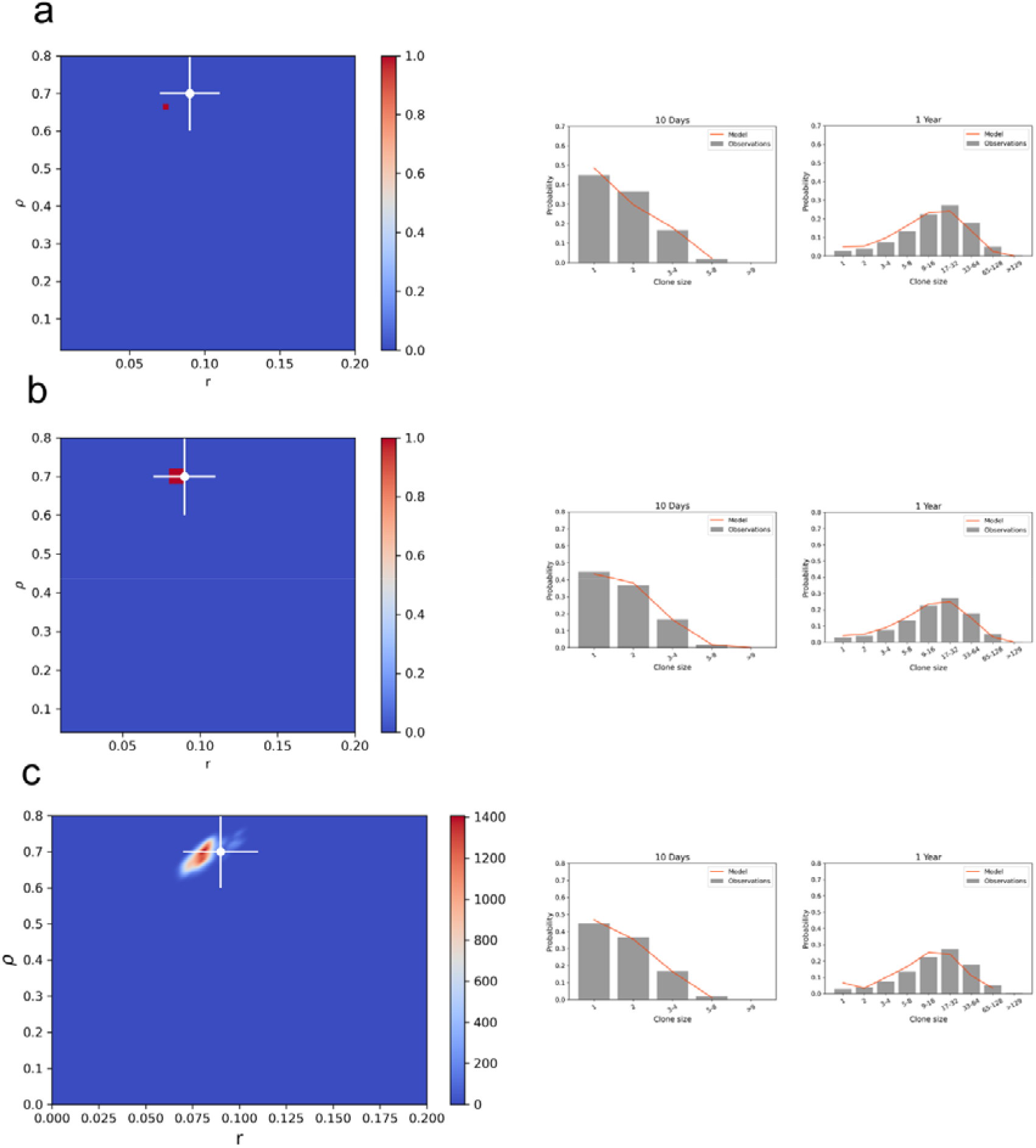
ABC approaches efficiently and appropriately parameterize the single progenitor model from lineage tracing data. a) Synthetic datasets where realistic cell cycle distribution times and biological variation are modelled show that likelihoods calculated from the analytical engine are overly conservative and inaccurate. Heatmap plots likelihood distribution calculated using the analytical solution. b) MLE based simulations considering Non-Markovian cell cycle distribution times improve parameter estimation but still fail to give smooth likelihood distributions. Heatmap plots likelihood distribution. c) An ABC approach to inferring parameters gives a smooth distribution and reasonable confidence intervals. Heatmap plots Kernel Density Estimate of final population of parameter sets. a-c: Input parameters and error estimates indicated by cross and error bars (2×SD). Synthetic datasets were generated using a mean r: 0.09 ± 0.01 SD, ρ: 0.7 ± 0.05 SD, λ: 2.9 ± 0.1 SD and assuming Gamma distributed cell cycle times. Right: Basal clone size probability distributions of the input synthetic data (grey) and the inferred parameters (orange) as obtained at an early and late timepoint.

In contrast to the analytical approach, simulation based techniques have the flexibility to account for more realistic cell cycle times. Simulation based MLE accurately calculated the input parameters and successfully reproduced the expected clone size distributions (Fig.3b, Supplementary Figure 3). However, the confidence intervals do not reflect the biological variation in the samples, substantially underestimating the uncertainty in *r* and *ρ* (Fig. 3b).

To address this, a Sequential Monte Carlo ABC (SMC-ABC) approach using a non-Markovian simulator was applied. We find that this generates a broad, smooth distribution that reflects the input parameters and uncertainty (Fig 3c, Supplementary Figure 3). Compared with results from MLE approaches, these findings demonstrate that the SMC-ABC inference technique is the most appropriate method for analysing lineage tracing datasets, whilst also being substantially more efficient.

### The contribution of individual timepoints to the likelihood distribution is sensitive to both the specific time and biological variation

Previous studies have used the results from individual timepoints to make arguments about the selection and rejection of different models (13). The underestimation of noise by MLE approaches raises the question of how biological variation influences the distributions observed at each individual timepoint. Related to this is the question of how each timepoint contributes to the wider likelihood. Answering these questions would aid experimental interpretation, confirming whether individual timepoints should be studied in isolation, but also offer a route to optimising experimental design.

To explore these questions, we revisited the experimental datasets and used synthetic datasets with modelled biological variation and calculated the likelihood distributions. As we are primarily interested in the shape of the distributions rather than the specific parameters proposed, we used the analytical MLE approach. The parameter likelihood distributions calculated from individual timepoints vary strongly across time and have extremely broad distributions, frequently covering large parts of parameter space. As such they are insufficient to estimate parameters in isolation (Fig. 4a). We further conclude that the precise estimation of likelihood from the whole timeseries effectively arises from the overlap of these distinct, broad distributions-that is to say, individual timepoints do not strongly point to a small distribution of parameters, but when considered together, they only agree on a narrow set.

**Fig. 4:**
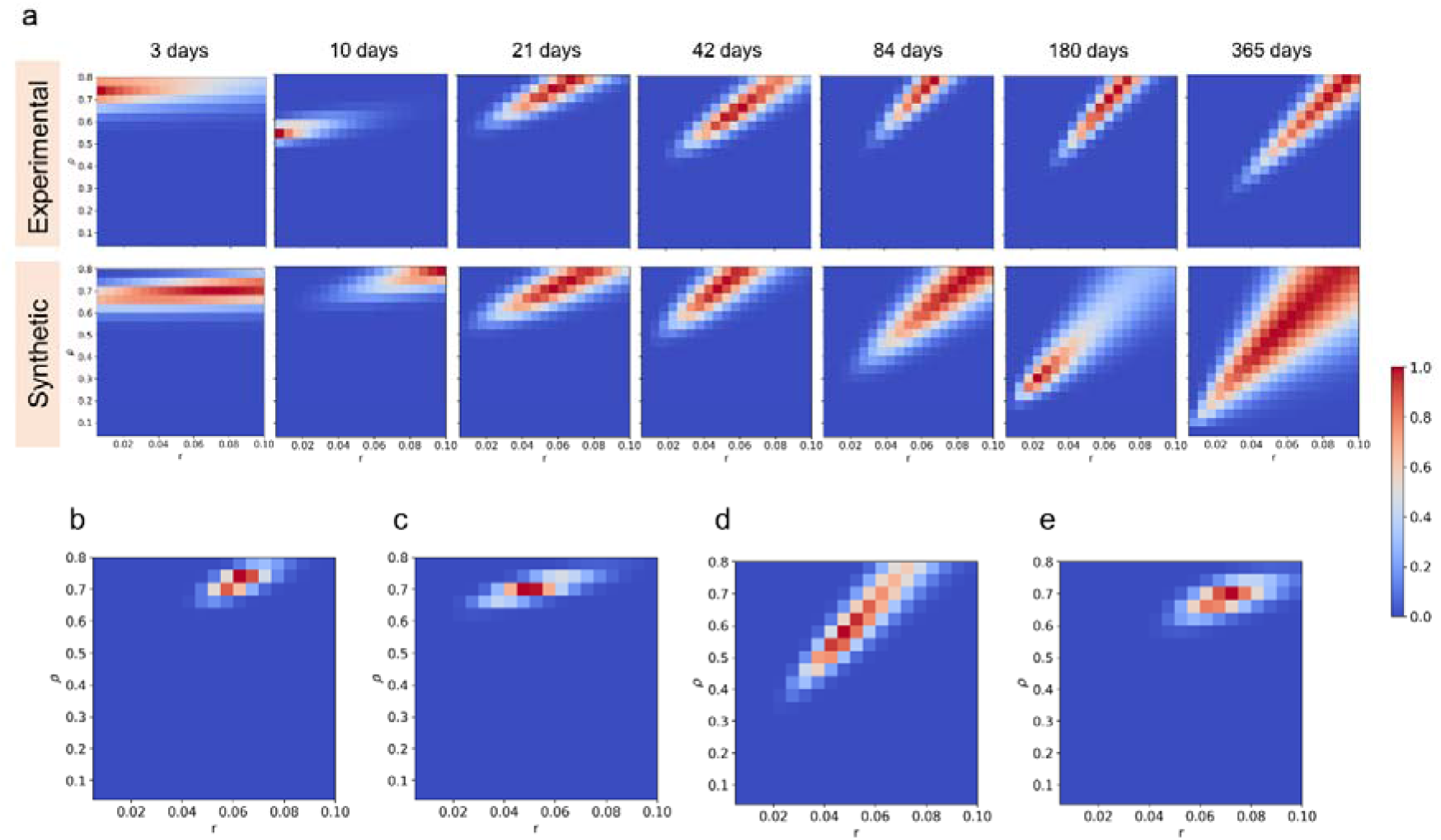
Estimated likelihood distributions across individual timepoints are highly sensitive to biological variation. a) Parameter likelihood distributions across individual timepoints calculated by the analytical engine on mouse oesophagus lineage tracing data from (3) (top) and synthetic datasets with biological variation (bottom). Likelihoods from individual timepoints are insufficient to estimate parameters on their own. b-e) Inferred parameter likelihoods estimated by the analytical engine on synthetic data with biological variation and increased sampling (5 mice per timepoint) considering all timepoints: 3, 10, 21, 42, 84, 180, 365 days (b), the three earliest timepoints: 3, 10, 21 days (c), the three latest timepoints: 84, 180, 365 days (d), a combination of early, middle and late timepoints: 3, 42, 365 days (e). a-e: Synthetic datasets were generated using a mean r: 0.08 ± 0.02 SD, ρ: 0.7 ± 0.1 SD, λ: 2.85 ± 0.15 SD and assuming exponentially distributed cell cycle times.

When compared with likelihood distributions calculated from experimental measurements, whilst we find that whilst the distributions are generally highly similar, the small number of biological replicates at each timepoint (two or three) leaves them prone to distortion by chance parameter combinations (Fig. 4a, 180 days). This would suggest that individual timepoints should not be used to draw strong conclusions as they are highly prone to chance variation.

One feature of note is that whilst the likelihood varies over time, later timepoints become increasingly similar. This raises the question of whether all timepoints need to be collected, and whether some timepoints could be omitted to increase sampling at other timepoints, potentially reducing the cost of the experiment and making individual timepoints more reliable. To investigate this, parameter likelihoods were calculated for synthetic datasets with increased sampling per timepoint (Fig. 4b). We then performed subsequent likelihood estimations considering different timepoint combinations (Fig. 4c-e), whilst maintaining the total number of synthetic “mice” considered. We find that including just early (Fig. 4c) or late (Fig. 4d) timepoints was insufficient to estimate the correct parameters of the synthetic datasets, a combination of one early, middle and late timepoint was able to estimate the expected likelihood distributions successfully (Fig. 4e).

## Discussion

Epithelial tissue maintenance is particularly important to understand homeostasis and processes such as aging, preneoplasia and cancer formation. The advent of genetic lineage tracing has provided useful insights on epithelial stem cell fate decision processes. Such studies are increasingly popular and this underlines the need for appropriate tools to analyze such datasets. Stem cell dynamics in multiple epithelial tissues has been argued over several years to be described by the single progenitor model (1–4,7), a mathematical model described with only three parameters, the division rate (*λ*), a stratification rate (*Γ*) and a probability of symmetric division (*r*). Despite this simplicity, we showed here how the choice of statistical approach can undermine the analysis, and how simulated datasets can be used to validate the approach taken. A key finding was that maximum likelihood calculations based on either an analytical approach (5) or simulation was overly precise, and produced unrealistic estimations of uncertainty that had the potential to obscure true parameters. In extremis, when tested on synthetic datasets that explicitly accounted for realistic cell cycle distributions and inter-mice biological variation, the proposed parameter values derived from the analytical solution were both inaccurate and the true values were outside confidence intervals. Simulation based maximum likelihood approaches were less inaccurate than the analytical solution, but had similar issues with uncertainty, and their utility was limited by their computational cost. In addition to the supercomputing power required for this analysis, the requirement to perform large numbers of simulations additionally limited the granularity of the analysis, effectively reducing the number of parameters that were tested. We found that despite being substantially less intensive, the SMC-ABC approach was able to account of realistic cell cycle times and identify the input parameters accurately with realistic error estimates. This tool, presented here and distributed with this manuscript, allows for accurate and efficient analysis of newly collected datasets following our protocol. It should be noted that whilst this tool enables the analysis of epithelial tissues, it can also be applied in other systems where cohesive clones are observed.

A major motivation for refining experimental design is to maximise the information generated whilst reducing costs and in particular reducing the need for animal use. Our findings suggest that increasing sampling at individual timepoints whilst reducing the total number of timepoints would increase the reliability of individual timepoints without impacting parameter estimation. Whilst this has implications on experimental design in transgenic systems, it also illustrates how simulation and modelling can aid experimental design beyond statistical tests. Here, simulating clone growth enables us to efficiently explore how parameters may be measured and the implications of well known confounders like inter-mouse variation on what can be safely interpreted from a dataset. The adoption of similar approaches can be both used before the experiment is run, to establish a protocol, but also afterwards to confirm whether unexpected features of the data can be trusted.

## Methods

### The single progenitor model

The single progenitor model supports the idea that the tissue is maintained by a single, equipotent progenitor population of basal cells that are able to give rise to either stem cells or differentiating daughters stochastically. The progenitor cell compartment in the basal layer is modelled as containing a mixture of stem cells, which go on to divide, and differentiated cells, which go on to stratify into upper layers of the tissue. Cell fate is determined on cell division; a stem cell divides to either produce one differentiated and one stem cell (asymmetric division), or either a pair of differentiated cells or a pair of stem cells (symmetric division) (Fig 1b). This population asymmetric mode of tissue renewal can be described as a continuous time Markovian process, as shown by (1–3) Eq. (1):

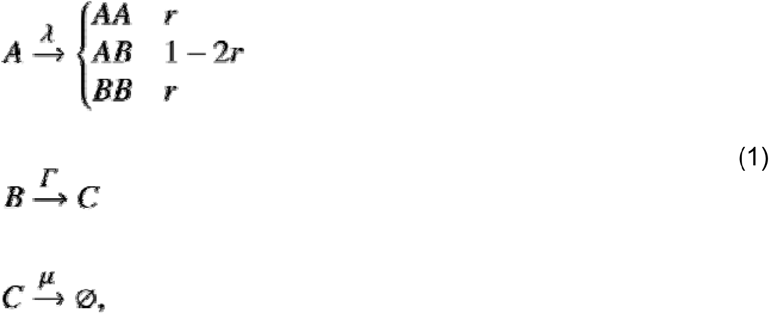

where represents the basal layer progenitor cells, the basal cells committed to differentiate and the suprabasal layer cells. Progenitor cells divide regularly with an overall division rate and give rise to either two progenitor daughters ( ), two differentiating daughters ( ) or one daughter of each type ( ) with fixed probabilities. Given the fact that ***AA*** symmetric division leads to clone expansion and symmetric division tends towards clone extinction, the two symmetric division rates should be equal in order for a steady state in terms of number of cells to be maintained across the progenitor clone population. The probabilities of symmetric and asymmetric divisions are and respectively with. Differentiating daughters in the basal layer stratify to the suprabasal layer at rate and supra basal cells,, are shed at rate. The fraction of cells in the basal layer that go on to divide is ρ. As the rules of homeostasis dictate that the total basal layer cell population stay constant over time, an additional rule arises, as shown by (1–3):

______

Therefore, parameterising the model requires specific values to be found for,, and.

### Generating clone size distributions following the single progenitor model

As the single progenitor model obeys a continuous time Markov process, the time evolution of proliferating (A) and differentiating (B) cell populations can be formulated in terms of the stochastic Master equation:

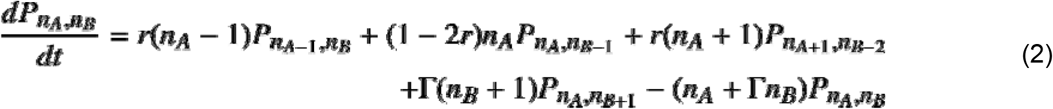

where denotes the probability of finding clones containing proliferating cells and differentiated cells. Antal and Krapivsky sought to provide an exact formula for the Master equation, calculating clone size probabilities,, and clone survival probabilities. The exact solution was obtained as shown in Eq. (3) (the full derivation is given in (5)).

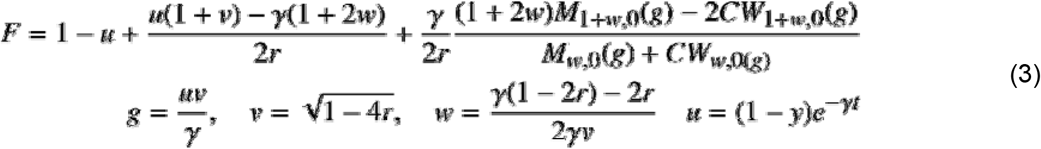

C is a constant determined as shown in Eq. (4):

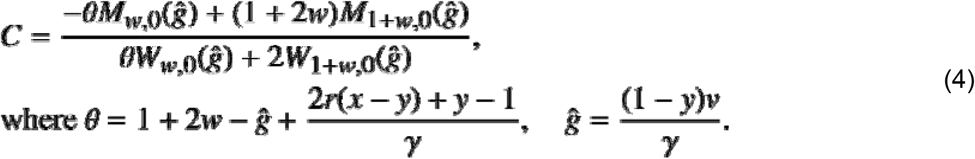

The analytical formula relies upon the use of confluent hypergeometric functions (Whittaker functions denoted by the terms *W* and *M*). To use the analytical solution in our analysis, we implemented the analytical formula in Python. Specifically, for every *r*, *ρ* combination (0 < *r* < 0.5 and 0 < *ρ* < 1) and for every time point Eq. (3) was called to compute the probability for a given basal clone size *n*. Clone size probabilities were searched for a set of 49×49 *r* and *ρ* parameter combinations. As *λ* was measured independently from H2BGFP dilution assays and provided to us, we used a fixed *λ* value when computing clone size probabilities for both experimental and synthetic datasets.

As an alternative to the analytical solution, the time evolution of clonal populations following the single progenitor paradigm was also simulated using the Gillespie stochastic simulation algorithm (14). Specifically, for a fixed *λ* value and for 19×19 *r*, *ρ* parameter combinations (0 < *r* < 0.5 and 0 < *ρ* < 1), multiple Gillespie simulation repetitions were performed (*N* = 100,000) to estimate the probability of observing a given basal clone size *n* at a given time point *t*. Initially single progenitor model simulations were performed assuming exponentially distributed cell cycle times. Non-Markovian simulations of the single progenitor model were performed assuming a Gamma-shaped cell cycle time distribution (6).

Synthetic datasets were generated by performing multiple Gillespie single progenitor model simulations under a specific *λ*, *r*, *ρ* parameter set. Synthetic datasets with biological variation and realistic cell cycle times were generated by performing simulations that took into account the number of mice typically used in a lineage tracing experiment (two to three per timepoint). Specifically, we performed multiple Non-Markovian single progenitor model simulations (*N* = 10,000), considering 7 different timepoints. This process was repeated 21 times, therefore simulating 21 animals so that 3 mice were considered per each time point. The simulation set corresponding to an individual mouse was assigned slightly different *λ*, *r*, *ρ* values drawn from a normal distribution to include inter-mice variability (*λ* = 2.9 ± 0.1, *r* = 0.09 ± 0.01, *ρ* = 0.7 ± 0.05).

### Single progenitor parameter inference

To fit the single progenitor model simulations against clonal data sets and identify appropriate parameter values, the following inference techniques were tested. Fitting was performed on parameters and.

#### Maximum Likelihood Estimation (MLE)

To calculate the likelihoods for the SP model parameters, a grid search was performed on a range of valid parameter values ( and ) and the theoretical estimates of basal clone size distributions - obtained either analytically or by performing stochastic Gillespie simulations – were contrasted with the ones observed experimentally by assessing the log-likelihood of every parameter combination. The most probable parameter combination was then selected as the parameter set with the maximum log-likelihood,

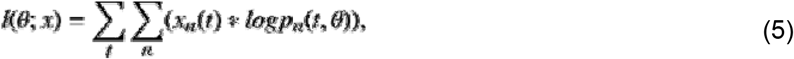

where corresponds to the frequency of measured clone sizes with n basal cells at time and is the probability of observing clones of size at time for a given parameter set values.

#### Sequential Monte Carlo Approximate Bayesian Computation (SMC-ABC)

To infer the parameters for the SP model an SMC-ABC approach was followed (9–15). Simulations of the SP model were performed starting from initial and values ( and ) drawn from a uniform distribution, used as prior. For every simulation round (population), a distance metric was computed for every value pair based on the sum of the Kolmogorov Smirnov (KS) test’s distance summary statistic.

While iterating over successive populations ( ), the new parameter sets to be tested were derived from a resampled and perturbed weighted set of points previously drawn. Perturbation allows a better exploration of the parameter space. Parameter values with calculated distance above a certain threshold (tolerance) were rejected, thus aiming to obtain the posterior distribution after certain rounds of tries. Tolerance is decreased after each round. The population size (number of particles to be accepted at each round) was set to 500.

The runtimes of the different analysis workflows were the following: The total CPU time for the analytical solution with MLE considering a 49×49 grid of parameters values was 10.2 hours. The total CPU time for the MLE based simulations considering 100,000 clone simulations per 19×19 combinations of parameters was 28 hours. The total CPU time for the SMC-ABC considering 10 populations, a population size of 500 and simulating 1000 clones per parameter combination was 38.33 hours. All analyses ran on a single core of an Intel(R) Xeon(R) processor (E5-2698 v3 @ 2.30 GHz).

All code was implemented in python 3.6, in Jupyter notebooks, using numpy. Sequential Monte Carlo ABC was used through the pyABC library. Code is available in supplementary information.

## Supporting information

Jupyter notebooks for analysis

## Acknowledgements

We thank Allon Klein, and the Hall and Jones groups for useful discussions.

## References

1. Clayton E, Doupé DP, Klein AM, Winton DJ, Simons BD, Jones PH. A single type of progenitor cell maintains normal epidermis. Nature. 2007;446(7132):185–9. Available from: http://www.ncbi.nlm.nih.gov/pubmed/17330052

2. Doupé DP, Klein AM, Simons BD, Jones PH. The Ordered Architecture of Murine Ear Epidermis is Maintained by Progenitor Cells with Random Fate. Dev Cell. 2010;18(2):317–23.

3. Doupé DP, Alcolea MP, Roshan A, Zhang G, Klein AM, Simons BD, et al. A single progenitor population switches behavior to maintain and repair esophageal epithelium. Science. 2012;337(6098):1091–3. Available from: http://www.ncbi.nlm.nih.gov/pubmed/22821983

4. Lim X, Tan SH, Koh WLC, Chau RMW, Yan KS, Kuo CJ, et al. Interfollicular epidermal stem cells self-renew via autocrine Wnt signaling. Science. 2013;342(6163):1226–30. Available from: http://www.ncbi.nlm.nih.gov/pubmed/24311688

5. Antal T, Krapivsky PL. Exact solution of a two-type branching process: Clone size distribution in cell division kinetics. J Stat Mech Theory Exp. 2010;P07028.

6. Piedrafita G, Kostiou V, Wabik A, Colom B, Fernandez-Antoran D, Herms A, et al. A single-progenitor model as the unifying paradigm of epidermal and esophageal epithelial maintenance in mice. Nat Commun. 2020 Dec 18;11(1).

7. Rompolas P, Mesa KR, Kawaguchi K, Park S, Gonzalez D, Brown S, et al. Spatiotemporal coordination of stem cell commitment during epidermal homeostasis. Science. 2016;9(6):855–61. Available from: http://www.ncbi.nlm.nih.gov/pubmed/27229141

8. Alcolea MP, Greulich P, Wabik A, Frede J, Simons BD, Jones PH. Differentiation imbalance in single oesophageal progenitor cells causes clonal immortalization and field change. Vol. 16, Nature cell biology. 2014. 615–22 p. Available from: http://dx.doi.org/10.1038/ncb2963

9. Toni T, Welch D, Strelkowa N, Ipsen A, Stumpf MPH. Approximate Bayesian computation scheme for parameter inference and model selection in dynamical systems. J R Soc Interface. 2009;6(31):187–202.

10. Toni T, Stumpf MPH. Simulation-based model selection for dynamical systems in systems and population biology. Bioinformatics. 2010 Jan 1;26(1):104–10. Available from: https://doi.org/10.1093/bioinformatics/btp619

11. Jain RB, Wang RY. Limitations of maximum likelihood estimation procedures when a majority of the observations are below the limit of detection. Anal Chem. 2008;80(12):4767–72.

12. Sugasawa S, Noma H. A unified method for improved inference in random effects meta-analysis. Biostatistics. 2019;1–29.

13. Mascré G, Dekoninck S, Drogat B, Youssef KK, Broheé S, Sotiropoulou P a, et al. Distinct contribution of stem and progenitor cells to epidermal maintenance. Nature. 2012;489(7415):257–62. Available from: http://www.ncbi.nlm.nih.gov/pubmed/22940863

14. Gillespie DT. Exact stochastic simulation of coupled chemical reactions. J Phys Chem. 1977 Dec;81(25):2340–61. Available from: http://pubs.acs.org/doi/abs/10.1021/j100540a008

15. Beaumont MA. Approximate Bayesian Computation in Evolution and Ecology. Annu Rev Ecol Evol Syst. 2010;41(1):379–406.

