## Supplementary Figures for "Methods for analysing lineage tracing datasets"

**Supplementary Figures and Legends**

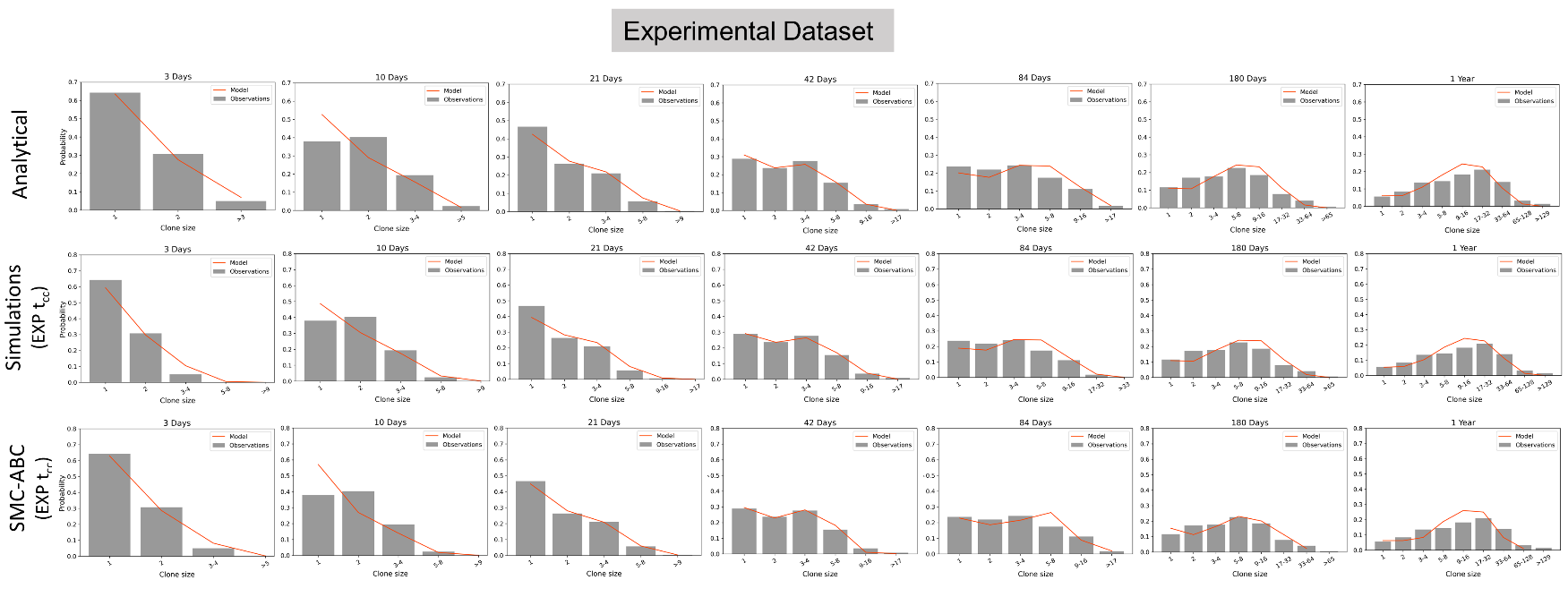

**Supplementary Figure 1:** **Observed vs simulated basal clone size probability distributions across all timepoints.** Observed and simulated basal clone size distributions are more dissimilar during early time points. Observed clone size distributions retrieved from clonal data provided in (1) and are depicted in grey bars. Orange lines show the modelled clone size distributions of the inferred most likely parameter combination as obtained by the three different analysis methods. All analyses were performed assuming an exponentially distributed cell cycle time.

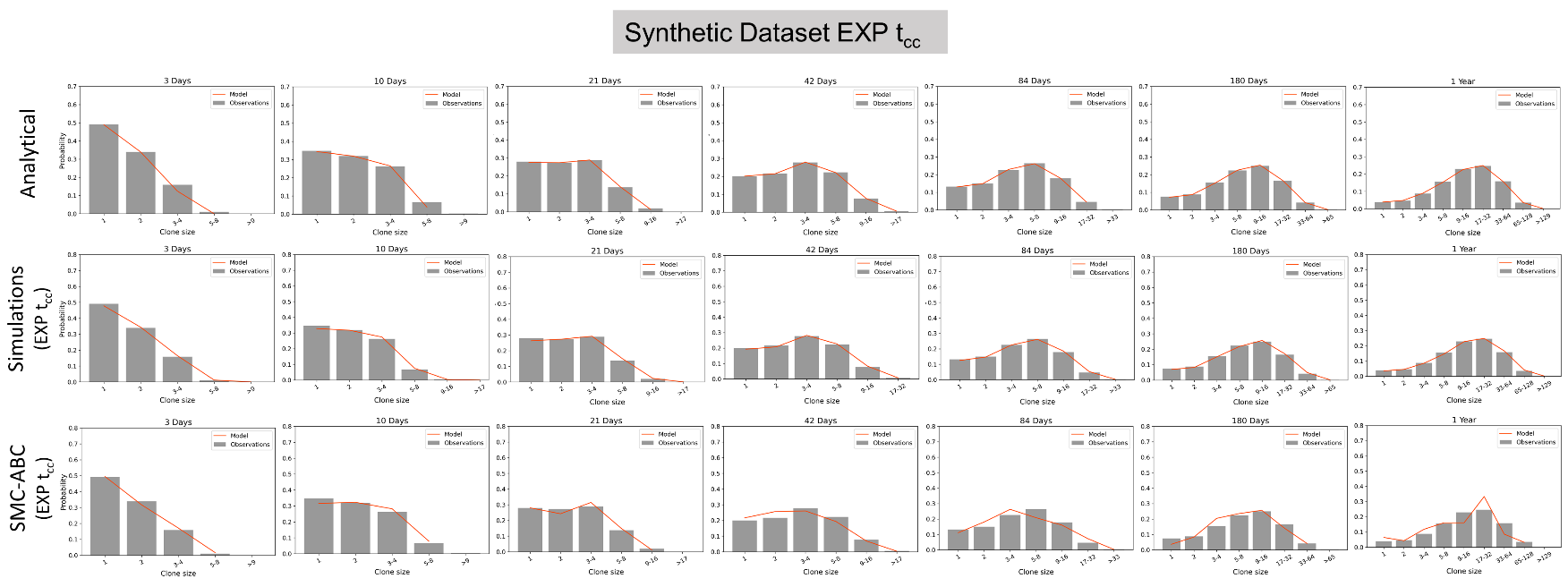

**Supplementary Figure 2:** **Observed (synthetic data) vs simulated basal clone size probability distributions across all timepoints.** An overall agreement between the observed and simulated basal clone size distributions was observed when synthetic data with no biological variation was analysed. Observed clone size distributions correspond to a synthetic dataset generated based on known $r, \rho$ parameter values $(r=0.06, \rho=0.5)$ and are depicted in grey bars. Orange lines show the modelled clone size distributions of the inferred most likely parameter combination as obtained by the three different analysis methods. All analyses were performed assuming an exponentially distributed cell cycle time.

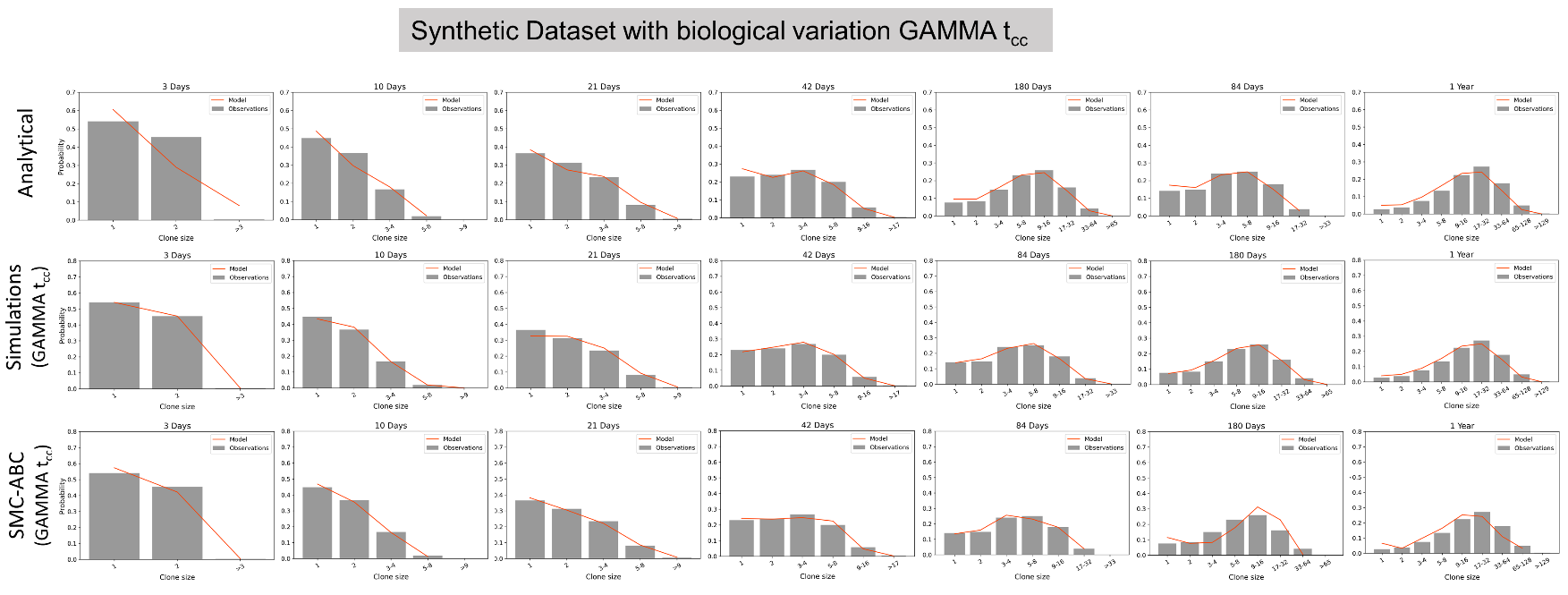

**Supplementary Figure 3:** **Observed (synthetic data with biological variation) vs simulated basal clone size probability distributions across all timepoints.** When more realistic datasets are considered, the analytical approach produces more dissimilar clone size distributions at the early time points. Observed clone size distributions (grey bars) correspond to the synthetic data generated using a mean $r: 0.09 \pm0.01 SD, \rho: 0.7 \pm0.05 SD, \lambda: 2.9 \pm0.1 SD$ and assuming Gamma distributed cell cycle times. Orange lines show the modelled clone size distributions of the inferred most likely parameter combination as obtained by the three different analysis methods. Simulation based MLE and the SMC-ABC analyses were performed assuming a Gamma distributed cell cycle time.

**References**

1.	Doupé DP, Alcolea MP, Roshan A, Zhang G, Klein AM, Simons BD, et al. A single progenitor population switches behavior to maintain and repair esophageal epithelium. Science. 2012;337(6098):1091–3. Available from: http://www.ncbi.nlm.nih.gov/pubmed/22821983
