## Supplementary material for "Methods for analysing lineage tracing datasets": Jupyter notebooks for analysis: MLE.ipynb

```
# %% [markdown]
## Please be aware that this is not the recommended workflow for analysis


# %% [markdown]
# Parameterising lineage tracing clonal data using MLE approaches

## Introduction

Over the past years, several studies based on lineage tracing became available, trying to analyse epithelial maintenance.
The single progenitor model has been shown to describe the observed stem cell dynamics in epithelial tissues accurately. Using three parameters, the division rate, the stratification rate, and the probability of symmetric division, this model predicts average clone size, clone size distributions, tissue homeostasis, and cell survival probabilities.

According to this model, the stem cell compartment is modelled as containing a mixture of stem cells, which go on to divide, and differentiated cells, which go on to stratify into upper layers of the tissue. Cell fate is determined on cell division; a stem cell divides to either give one differentiated and one stem cell daughters (asymmetric division), or either a pair of differentiated cells or a pair of stem cells (symmetric division). The probability of either symmetric division is r, and of asymmetric division is 1-2r. The model has two further parameters, the division rate (λ) and the stratification rate (Γ). The stratification rate can be expressed in terms of the division rate and the fraction of stem cells in the basal layer (ρ). In parameterizing the model therefore specific values are found for r, λ, and ρ. 

In this notebook we provide a workflow for analysing lineage tracing clonal datasets using either sigle progenitor model simulations or an analytical solution which both allow for maximum likelihood calculations.

<img src="Cartoon.png" alt="Drawing" style="width: 600px;"/>

# %% [markdown]
## Dependencies

In order to run the analysis provided by this notebook, the following libraries need to be installed:
mpmath
numpy
pandas
matplotlib 
scipy

Below we provide the option for automatically install the required libraries. Please bear in mind that this might affect other Python workflows running in your system. If you decide to use this option please convert the following block from Markdown to Code:

# %% [markdown]
import sys
(!sys.executable) -m pip install mpmath numpy pandas matplotlib scipy

# %% [markdown]
## User configuration

The following variables allow the user to choose their preferred analysis approach through the *analysis_plan* dictionary. Then the user can provide their input data and configure analysis parameters.

The following analysis options are available:
1. method: **'simulation'** or **analytical**.
    In the *'simulation'* mode, 'simulations of the single progenitor model are performed to estimate the probabilities of clone sizes at a given time point under different r,ρ parameter combinations. In the *'analytical'* mode clone size probabilities are computed by using a published analytical formula. 
2. individual_timepoints **'True'** or **False**.
    Clone size theoretical estimates are compared with the experimetally counted clone sizes using Maximum Likelihood estimation (MLE). In the *'False'* mode, the most probable parameter combination is selected as the one with the maximum log-likelihood across all time points. In the *'True'* mode MLE calculation is separated across individual time points.

# %% [code]
analysis_plan = {
    'method': 'analytical', # choose between 'simulation' and 'analytical'
    'individual_timepoints': False, # True if you want to perform the analysis across individual time points
}

## Path to input clonal data (please see below for correct formatting)
inputData = "./exampleClonalData.tsv"
# Average division time inferred experimentally
meanDivisionTimeInDays = 2.4

# Max number of generations for a given simulated clone (for simulations only)
generations = 230
# Number of cell clones to be simulated (for simulations only)
sampleSize = 100000
# specify how many parameter combinations will be searched
parameterBins = 20

# set r and rho ranges to search
max_rho = 0.8 # specify maximum rho value to be searched (0 < rho < 1)
max_r = 0.2 # specify maximum r value to be searched (0 < r < 0.5)
rhoRange = [(i*max_rho/parameterBins) for i in range(1,parameterBins)]
rRange = [(i*max_r/parameterBins) for i in range(1, parameterBins)]

warningSuppression = True # Suppresses warning messages notifying about pathological values (i.e. out of bound probabilities) in the analytical workflow 

# %% [markdown]
Please provide your input clonal data according to the following format:

Format for input clonal data file (tab delimited)<br />
First row: time points<br />
First column: clone size<br />
Rest: number of clones

# %% [raw]
	3	10	21	42	84	180	365
1	90	96	140	73	83	40	12
2	43	102	79	60	77	59	18
3	4	34	38	44	38	43	15
4	3	15	25	26	47	18	14
5	0	4	11	22	22	27	6
6	0	2	4	10	18	23	11
7	0	0	1	3	14	16	8
8	0	0	1	4	7	12	6
9	0	0	0	0	8	14	9

# %% [markdown]
## Load modules

Import necessary modules. The following libraries must be imported in order for the workflow to run successfully.

# %% [code]
%matplotlib inline

import os
import tempfile
import cmath
import math
import random
from functools import partial

from mpmath import *
import numpy as np
import pandas as pd
from scipy import stats
import matplotlib.pyplot as plt
import scipy as sp


# %% [markdown]
# Simulating cell fates

<img src="CellFate.png" alt="Drawing" style="width: 600px;"/>

The first behaviour we will simulate is fate determination of cells in a clone, starting from a single cell. This is the pattern of changing cell states as cells go on to divide. Examples of cell fates are drawn as dendrograms above. In (A), A cell divides, but neither daughter cell is a stem cell and so division stops. In (B), the divisions form a more complex (and unfinished pattern)- in the first division the cell fates are asymmetric giving one stem cell and one differentiated cell. That stem cell divides to give two stem cells, and so on until either all daughter cells in the clone divide becoming differentiated cells, or we stop the experiment

In this specific situation we are only simulating the fates- that is to say, we know that division and stratification occur at different rates, but we are not taking account of that yet- we just record the order in which things happened in discrete time. Even without precise timings we expect the functions to give us some of the key properties of the clone growth. 

We define a clone through a record type, where we record when the clone was born (*birthtime*), when the cell left the clone (*lifetime*), the random number generator used to determine the next fate (*rng*), whether the cell is a stem cell (*stem*) and the probability of dividing to give daughters with the same cell fate (*r*). We also include a list of other clones that the clone has given birth to (*siblings*).

The function *simulate* simulates the life of a single cell. Given a clone record, a maximum number of divisions (*limit*), and a reference variable used to track unique identities (*nameMarker*) it loops through events in the cells life, giving a final clone with some variables changed. 

In each loop simulate first tests whether the cell has been simulated enough times, and if it has returning a modified version of the last state of the cell. This is either the last state with the lifetime set to the current time (and a Boolean variable, *survivor*, set to true to indicate this), or if the cell is not a stem cell at the end of the simulation, the last state with the lifetime set to the current time+1. 

Next, if the cell is still a stem cell, it creates a new daughter cell as a copy of the parent, with a new unique identity value. We then determine using r and a random number whether the daughter and parent are stem cells after division, defining a new cell to replace the parent and the daughter. Finally, we restart the loop using the replacement cell with the daughter added to the list of siblings.

This however is not sufficient- we want to simulate all the daugter cells as well to see how the clone grows. To do this we define a recursive function *corecloneSimulateComplete*, which simulates a single cell, then recursively runs on all of the daughter cells (and their daughter cells and so on), appending only the key information as a tuple to list reference. This function itself is run through *cloneSimulateComplete*, which sorts the final list by unique identifier (this is important for later functions).

Finally, the function *experimentDiscreteTime* is defined which takes a list of discrete timepoints, runs cloneSimulateDiscreteTime, and asks how many cells are alive at each time.

# %% [code]
'''
A cell should update to divide and die. As we can only count living cells, 
we only need to measure the lifetime of a cell or experiment, and store its 
siblings.

For the whole clone size, each of the siblings need to be simulated to their
stratification or the end of the experiment.

The cell updates in discrete time, with an aim that randomly generated times 
are projected back on a cohort a posteri to reduce calculation times.
'''
        
class Cell:
    def __init__(self,init,stem,r,rng):
        self.currentTime = init
        self.stem = stem
        self.r = r
        self.rng = rng
        self.lifetime = 0
        self.birthtime = self.currentTime
        self.survivor = False
        self.siblings = []
        
    def simulate(self,limit):
        cont = True
        while(cont):
            if self.lifetime > 0:
                cont = False
            elif self.currentTime>limit:
                #End experiment- lifetime set to currenttime+1 if not a stem cell, currenttime if stem with surv bool set
                #Infinity is inappropriate as its a float, so setting lifetime to current time
                #If a clone has infinity *before* the last experimental timepoint, the simulation needs extension
                if self.stem:
                    self.survivor = True
                    self.lifetime = self.currentTime
                else:
                    self.lifetime = self.currentTime+1
                cont = False
            else:
                if self.stem:
                    self.currentTime += 1
                    sibling = Cell(self.currentTime,self.stem,self.r,self.rng)
                    rand = self.rng.random()
                    if rand < self.r:
                        pass
                    elif rand < (self.r*2):
                        self.stem = False
                        sibling.stem = False
                    else:
                        sibling.stem = False
                    self.siblings.append(sibling)
                else:
                    self.currentTime += 1
                    self.lifetime = self.currentTime
    #rand < r: p = r, two stem cells are produced
    #r < rand < 2r: p = r, two differentiated cells are produced
    #2r < rand <1: p = 1-2r, one stem cell and one differentiated cells are produced

    
'''
Simulates a cell up to an event limit, storing the discrete times only
'''
def cloneSimulateDiscreteTime(i,limit,acc):
    i.simulate(limit)
    acc.append((i.birthtime,i.lifetime))
    for s in i.siblings:
        cloneSimulateDiscreteTime(s,limit,acc)
    return(acc)

'''
Simulates a cell up to an event limit, storing the cell as result
'''
def corecloneSimulateComplete(i,acc,parent,nameMarker,limit):
        i.simulate(limit)
        nameMarker[0]+=1
        name = nameMarker[0]
        acc.append((i.birthtime,i.lifetime,parent,i.survivor))
        for s in i.siblings:
            corecloneSimulateComplete(s,acc,name,nameMarker,limit)
        return((acc))
def cloneSimulateComplete(i,limit):
    return(corecloneSimulateComplete(i,[],-1,[-1],limit))

def experimentDiscreteTime(c,timePoints):
    limit = max(timePoints)
    lifespans = cloneSimulateDiscreteTime(c,limit,[])
    def within(a,b,c):
        if c>a and c<=b:
            return(1)
        else:
            return(0)
    return([sum([within(b,d,t) for (b,d) in lifespans]) for t in timePoints])
    

def printClone(c):
    def core(c,d):
        print("Gen\t%d" % d)
        c.printHistory()
        for i in c.siblings:
            core(i,d+1)
    core(c,0)

def seedExperiment(i):
    random.seed(i)
    return(experimentDiscreteTime(Cell(0,True,0.25,random),[2,4,8,10,30,50,100]))

def seedClone(i):
    random.seed(i)
    return(cloneSimulateComplete(Cell(0,True,0.25,random),100))

# %% [markdown]
# Converting clone lifespan to realistic times

The fate simulator is able to generate the orders of events in a clones lifetime, but in order to compare this with experimental data we need to use the outputs from that function, and transform them into floating point numbers that reflect the division and stratification rates. The most common way to do this is to draw the division times from an exponential distribution characterised by its mean, but we also know that the exponential distribution is unrealistic for cell division times. Here we explore two alternative distributions used in the literature; a short, deterministic delay (the delay or refractory period) followed by an exponential distribution, and a deterministic lag period followed by times selected from a gamma distribution (with an additional shape parameter). As the exponential distribution is a special case of the gamma distribution where the shape is 1., we can focus on using just the gamma distribution when we start to compare with experimental data.

We first define functions that allow us to set create a random number generator for a given distribution with a specific set of parameters (using the numpy library to generate the underlying distribution). In each case the overall mean time for events to occur and a random number generator is passed to the function. In addition to this, a delay may be passed (either as an absolute delay, or expressed as a fraction of the mean) and a shape parameter. In each case the object returned is a function that takes unit as an input and returns a new floating point number from the distribution.

In each case we use an object that preallocates a large number of random numbers (*total*) and defines functions that return one, or an array of random numbers of a specified size, cumulatively summed. The functions that return values determine whether the store of numbers will run out and regenerates a set periodically. This is written like this for performance purposes. 

# %% [code]
class TimeGenerate:
    def __init__(self, seed, params):
        np.random.seed(seed)
        self.total = 10000000
        self.params = params
        self.validate()
        # Generate "total" times
        self.regenerate()

    def regenerate(self):
        self.precalculated = np.ones(self.total)
        self.position = 0

    def next(self, n=1):
        if n + self.position > self.total:
            self.regenerate()
            return self.next(n)
        else:
            result = np.cumsum(
                self.precalculated[self.position:self.position + n])
            self.position += n
            return result

    def nextOne(self):
        if 1 + self.position > self.total:
            self.regenerate()
            return self.nextOne()
        else:
            result = self.precalculated[self.position]
            self.position += 1
            return (result)

    def nextTerminated(self, n=1, terminator=None):
        if n + self.position > self.total:
            self.regenerate()
            return self.next(n)
        else:
            times = self.precalculated[self.position:self.position + n]
            # This replaces a value! it will not be reused and eventually will
            # be regenerated though
            times[-1] = terminator.nextOne()
            result = np.cumsum(times)
            self.position += n
            return result

    def validate(self):
        pass

    def drawTimes(self, limit):
        counter = 0
        totalTime = 0
        while totalTime < limit:
            totalTime += self.precalculated[self.position + counter]
            counter += 1
            if counter + self.position > self.total:
                self.regenerate()
                return self.drawTimes(limit)
        return self.next(counter)


class HalfTime(TimeGenerate):
    def regenerate(self):
        self.precalculated = np.ones(self.total) / 2
        self.position = 0


class ExponentialGenerate(TimeGenerate):
    def regenerate(self):
        (mean) = self.params
        self.precalculated = np.random.exponential(mean, self.total)
        # draw samples from an exponential distribution
        self.position = 0


class UniformGenerate(TimeGenerate):
    def regenerate(self):
        self.precalculated = np.random.uniform(0, 1, self.total)
        # draw samples from a uniform distribution
        self.position = 0


class ExponentialGenerateDelay(TimeGenerate):
    def regenerate(self):
        (mean, delay) = self.params
        self.precalculated = np.random.exponential((mean - delay),
                                                   self.total) + delay
        self.position = 0

    def validate(self):
        assert (self.params[0] > self.params[1])


class GammaGenerate(TimeGenerate):
    def regenerate(self):
        (shape, scale) = self.params
        self.precalculated = np.random.gamma(shape, scale, self.total)
        # draw samples from a gamma distribution
        self.position = 0


class GammaGenerateDelay(TimeGenerate):
    def regenerate(self):
        (mean, shape, delay) = self.params
        scale = (mean - delay) / shape
        self.precalculated = np.random.gamma(shape, scale, self.total) + delay
        self.position = 0

    def validate(self):
        assert (self.params[0] > self.params[2])


# More elegant way of expressing the delay
class GammaGenerateFractionalDelay(TimeGenerate):
    def regenerate(self):
        (mean, shape, delay) = self.params
        scale = (mean * (1 - delay)) / shape
        self.precalculated = np.random.gamma(shape, scale,
                                             self.total) + delay * mean
        self.position = 0

    def validate(self):
        assert (0 <= self.params[2])
        assert (self.params[2] <= 1)

# %% [markdown]
## Converting discrete time to real time

The function *correctTime* below takes two random number generator objects (of the type we have just defined above), an initial time, and an array of tuples generated by the cloneSimulate function defined above. It returns a new array with the time of birth and the time of loss of the cell. 

To do this we first generate an array with random numbers drawn from the function representing the division times. If the cell stratifies we replace the last number in the array with a value drawn from the function representing stratification times. Finally, we perform a cumulative sum so that all elements represent the times taken for each event to happen from an initial timepoint (here arbitrarily set to zero).

This gives us a series of timepoints that are accurate relative to the time when the cell was first born. However, we need the times to be relative to the start of the experiment to be useful. To do this for each cell we need to look at the times of its parent (and in turn, its parents and grandparents until the original cell) and adjust all the timepoints by the time the cell was born. Uniquely, the original cell has a parent set to a negative value and if this is found in the loop, the initial time is set to the value passed when calling the function.

As an example, we want to generate times for the third cell in a clone. The third cell is a daughter of the first. The discrete times are 

cell 1 born @ 0, lost @ 3
cell 3 born @ 1, lost @ 2

This will give us initially two arrays of timepoints;

cell1 = [|0.;0.5;1.0;1.5;|]
cell3 = [|0.;1.0|]

For cell 3, assuming that the initial time of the experiment was 0 (so cell 1 remains unchanged), we then need to add 0.5 (the first time from cell 1) to every timepoint. We get this by using the discrete birthtime as an index in the array. Finally, we return the first and last elements of the new array as the time of birth and the time of death; 0.5 and 1.5.

This relies on being able to look up the parent time. Earlier we sorted the array returned by cloneSimulate by the unique identifier- this allows us to use the parent id (stored in the tuple) as an index in the original search. It also ensures that we correct the times in order of cell creation- this is important as when we correct the times we mutate the values in the original array, and ensures that we have always corrected the parents before we update their children. 

We now have enough information to directly compare our results with lineage tracing data.

# %% [code]
def correctTime(c,initial,division,stratification,limit):
    #c is tuple of birth,death,parent,survival (int*int*int*bool)
    #First get all the relative times for all the clones, measured from birth. 
    eventTimes = [((division.next(d-b),b,d,p) if s else (division.nextTerminated(d-b,terminator=stratification),b,d,p) ) for (b,d,p,s) in c ]
    
    #Second correct the intial times of all of the events and store to the acc
    def includeBirth(times,birthday,deathday,parent):
        #modifier is going to be calculated from the birthtime, and the time of the parent
        #this relies on mutables. Potential for ugliness
        #Your parent's life times are stored by index- so if you parent is x, the lifetimes are in eventTimes[x]
        #Times are in the first element of the eventTimes tuple- eventTimes[x][0]
        #Your birthday (n) is the (n-1)th event in the parents life- so its in eventTimes[x][0][n-1*]
        #Your parents birthday needs to be taken into account too; so if your parent was born on 3 and you on 6 you need n-pbday-1
        #print(times)
        if parent<0:
            #special case- the initial cell
            times+=initial
            return((initial,times[-1])) 
        else:
            parentBirthday = eventTimes[parent][1]
            timeOnBirth=eventTimes[parent][0][birthday-parentBirthday-1]
            if(len(times)):
                times+=timeOnBirth
                return( (timeOnBirth, times[-1] ) )
            else:
                #special case- this is a cell born on the last time point. Arbitrarily set to infinity
                #Note differentiated cells born on the last timepoint have their lifetime extended, so this is only stem cells
                return((timeOnBirth,math.inf))
    acc = [includeBirth(times,birthday,deathday,parent) for (times,birthday,deathday,parent) in eventTimes]
    
    return(acc)
    # For examle, gammagenerator will generate s sequence of random values, a,b,c,d,e,f,g,,,
    # the original cell (O) divides into two daughtor cells (A0 and A1), and A0 divides into B0 and B1
    # the length of lifetime for cellO would be a, and for A0 (a+b0), A1 (a+b1), B0 (a+b0+c0), B1 (a+b0+c1),,,
    # the time point of bithtime and deathtime are then calculated based on the birthtime and lifetime of the parent
    # i.e. cumulative sum

# %% [markdown]
# Analytical solution for clone size probabilities

Here we provide the necessary functions of a published analytical solution for calculating clone size probabilities as provided in Antal, T. & Krapivsky, P.L. Exact solution of a two-type branching process: clone size distribution in cell division kinetics. *J Stat Mech Theory Exp* P07028 (2010). https://iopscience.iop.org/article/10.1088/1742-5468/2010/07/P07028 

The published analytical formula is implemented in function *F*. Functions *Psurv* and *Pn* compute clone survival and clone size probabilities respectively.

# %% [code]
def normal_round(n):
    if n - math.floor(n) < 0.5:
        return math.floor(n)
    return math.ceil(n)


def F(x, y, t, r, gamma):
    v = (1 - 4 * r) ** 0.5
    w = (gamma * (1 - 2 * r) - 2 * r) / (2 * gamma * (1 - 4 * r) ** 0.5)
    gg = (1 - 4 * r) ** 0.5 / gamma
    u = (1 - y) * np.exp(-gamma * t)
    u0 = (1 - y)
    Q = 1 + 2 * w - gg * u0 + (2 * r * (x - y) + y - 1) / gamma
    C = (-Q * whitm(w, 0, gg * u0) + (1 + 2 * w) * whitm(1 + w, 0, gg * u0)) / \
        (Q * whitw(w, 0, gg * u0) + 2 * whitw(1 + w, 0, gg * u0))
    f = 1 - u + \
        (u * (1 + v) - gamma * (1 + 2 * w)) / (2 * r) + \
        (gamma / (2 * r)) * ((1 + 2 * w) * whitm(1 + w, 0, u * gg) -
                             2 * C * whitw(1 + w, 0, u * gg)) / \
        (whitm(w, 0, u * gg) + C * whitw(w, 0, u * gg))
    return f


def Antal(z, t, r, gamma):
    g = []
    for k in range(len(z)):
        if z[k] == 1:
            g.append(1)
        else:
            g.append(F(z[k], z[k], t, r, gamma))

    return g


def Psurv(t, l, r, rho):
    rho = rho - 0.00001
    gamma = rho / (1 - rho)
    T = l * t
    p = 1 - np.real(Antal([0], T, r, gamma))
    if p <= 0 or p > 1:
        p = Psurv(t, l, r + 0.00001, rho - 0.001)
    return p


def Pn(n, t, l, r, rho, N, returnConsecutive='', max_n=''):
    if max(n) >= N:
        N = max(n) + 1
    rho = rho - 0.00001
    assert (rho < 1)
    assert (rho > 0)
    gamma = rho / (1 - rho)
    T = l * t
    Gvals = Antal([cmath.exp(2 * np.pi * 1j * k / N) for k in range(int(N))], T,
                  r, gamma)
    p = []

    if returnConsecutive:
        p = np.zeros(max_n)
        for lp in range(len(n)):
            temporary = [cmath.exp(-2 * np.pi * 1j * k * n[lp] / N) for k in
                         range(int(N))]
            p[n[lp] - 1] = np.real(
                (1 / N) * sum([a * b for a, b in zip(Gvals, temporary)]))
    else:
        for lp in range(len(n)):
            Temporary = [cmath.exp(-2 * np.pi * 1j * k * n[lp] / N) for k in
                         range(int(N))]
            p[lp] = np.real(
                (1 / N) * sum(a * b for a, b in zip(Gvals, Temporary)))
    tst = sum(p)
    if tst > 1 or tst < 0 or np.isnan(tst):
        print('Pathological point: r=' + str(r) + ' rho=' + str(
            rho) + '. Making 0.1% perturbation')
        p = Pn(n, t, l, r + 0.00001, rho - 0.001, N, returnConsecutive, max_n)

    return p

# %% [markdown]
## Import experimental data
Here we will parse the input file of clone counts and extract the necessary information.

# %% [code]
def getLineage(filename):
    #reads a tsv file of clone counts
    df = pd.read_csv(filename, sep="\t")
    counts = np.transpose(df.values).astype(float)
    header = df.columns.values
    return(header[1:].astype(float),counts[1:])

times, counts = getLineage(inputData)

#Reproduce the linear increase property observed in average clone size 
averageClone = np.array([sum([(size+1)*number for (size,number) in enumerate(t)])/sum(t)  for t in counts])
plt.plot(times,averageClone)

# %% [markdown]
## Comparing models with experiments using MLE
Here we provide a list of functions necessary for calculating parameter likelihoods based on the input clone count data.

To fit model to experimets the following steps are followed: 

1. For a range of valid r, ρ values (defined by the user in *rRange* and *rhoRange* in User configuration), a probability of observing a given clone size at a given time point is calculated from either a simulation approach or the analytical solution. The analysis type is also specified by the user. 
2. The experimental data is used to assess the log-likelihood of every parameter combination by comparing the observations (number of clones of a given size at a given time point) with the probabilities generated at the previous step. The most probable parameter combination is then selected using maximum likelihood estimation (MLE). If the user selects the *individual_timepoints* analysis, the MLE is calculated per each time point individually.

# %% [code]
def logLikelihood(obs, lp, cloneSizeBins=[]):
    maxClone = len(lp)

    if len(cloneSizeBins) > 0:
        obsBins = [sum(obs[sum(cloneSizeBins[0:n]):sum(cloneSizeBins[0:(n + 1)])]) for n in
                   range(len(cloneSizeBins))]
        return sum([obsBins[j] * lp[j] for j in range(maxClone)])
    else:
        return sum([obs[j] * lp[j] for j in range(maxClone)])


def sizesToFreq(sizes, cloneSizeBins):
    biggest = max([max(c) for c in sizes])
    sizeList = [[t.count(s + 1) for s in range(biggest)] for t in sizes]
    countPerTimePoint = [sum(s) for s in sizeList]
    binSizeList = [[sum(t[sum(cloneSizeBins[0:n]):sum(cloneSizeBins[0:(n + 1)])]) for n in
                    range(len(cloneSizeBins))] for t in sizeList]
    logFreq = [[math.log10(
        1 / 100000) if total == 0 or item == 0 else math.log10(item / total) for
                item in n] for n, total in zip(binSizeList, countPerTimePoint)]
    return logFreq


def fateToLogProbability(timePoints, fates, generations, div, strat, cloneSizeBins):
    times = np.asarray(
        [np.asarray(correctTime(c, 0, div, strat, generations)) for c in fates])

    def within(birth, death, time):
        if birth <= time < death:
            return 1
        else:
            return 0

    cloneSizes = [
        [sum([within(cell[0], cell[1], t) for cell in clone]) for clone in
         times] for t in timePoints]
    freq = sizesToFreq(cloneSizes, cloneSizeBins)
    return freq


def parameterLikelihood(rho, meanDivTime, timePoints, fates, generations, div, seed,
                  data, cloneSizeBins):
    stratTime = (1 - rho) * meanDivTime / rho
    strat = ExponentialGenerate(seed, stratTime)
    # Now get the probabilities
    lpBins = fateToLogProbability(timePoints, fates, generations, div, strat, cloneSizeBins)
    # Now get the likelihoods
    if analysis_plan['individual_timepoints']:
        likelihood = [logLikelihood(data[t], lpBins[t], cloneSizeBins) for t in
                range(len(data))]
    else:
        likelihood = sum([logLikelihood(o, p, cloneSizeBins) for o, p in zip(data, lpBins)])
    return likelihood


def searchRho(timePoints, r, rhoRange, sampleSize, seed, div, meanDivTime, data,
              cloneSizeBins):
    random.seed(seed)
    generations = 230

    def seedWrapper(r, generations, seed):
        random.seed(seed)
        clone = Cell(0, True, r, random)
        return (cloneSimulateComplete(clone, generations))

    fates = np.asarray(
        [np.asarray(seedWrapper(r, generations, i)) for i in range(sampleSize)])

    return ([
        parameterLikelihood(rho, meanDivTime, timePoints, fates, generations, div,
                      rhoSeed, data, cloneSizeBins) for rhoSeed, rho in
        enumerate(rhoRange)])


def calculateSimulationLikelihood(div, meanDivTime, data, timePoints, cloneSizeBins):
    result = np.array([searchRho(timePoints, r, rhoRange, sampleSize, seed, div,
                                 meanDivTime, data, cloneSizeBins) for (seed, r) in
                       enumerate(rRange)])
    return result

#### The following are specific to the analytical approach ####

def calculateLikelihood(probability, data):
    if analysis_plan['individual_timepoints']:
        lpBins = [math.log10(1 / 1000000) if n <= 0 else math.log10(n) for n in
                  probability]
        likelihood = logLikelihood(data, lpBins)
    else:
        # like = allLogLike(data, lpBins)
        lpBins=[[math.log10(1 / 1000000) if n <= 0 else math.log10(n) for n in t] for t in probability]
        likelihood = sum([logLikelihood(o, p) for o, p in zip(data, lpBins)])
    return likelihood


def calculateAnalyticalLikelihood(t, nRange, lambdaRange, rhoRange, rRange, data):
    maxN = max(list(map(max, nRange)))

    PScanPP = [[[[np.zeros(maxN) for L4 in range(len(nRange))] for L3 in
                 range(len(rRange))] for L2 in range(len(rhoRange))] for L1 in
               range(len(lambdaRange))]

    PSurvScanPP = [
        [[np.zeros(len(nRange)) for L3 in range(len(rRange))] for L2 in
         range(len(rhoRange))] for L1 in range(len(lambdaRange))]

    timePoints = range(len(t))
    nBadValues = 0

    for lambdaIterator in range(len(lambdaRange)):
        PScanPP_local = [
            [[np.zeros(maxN) for L4 in range(len(nRange))] for L3 in
             range(len(rRange))] for L2 in range(len(rhoRange))]

        PSurvScanPP_local = [
            [np.zeros(len(nRange)) for L3 in range(len(rRange))] for L2 in
            range(len(rhoRange))]

        for rhoIterator in range(len(rhoRange)):
            for rIterator in range(len(rRange)):
                for timePointIterator in timePoints:
                    probS = Psurv(t[timePointIterator], lambdaRange[lambdaIterator], rRange[rIterator],
                                  rhoRange[rhoIterator])
                    avgN = 1 / probS
                    Niter = max([normal_round(int(4 * avgN)),
                                 normal_round(1.5 * max(nRange[timePointIterator])), 10])
                    probN = Pn(nRange[timePointIterator], t[timePointIterator], lambdaRange[lambdaIterator], rRange[rIterator],
                               rhoRange[rhoIterator], Niter, 'returnConsecutive', maxN)

                    if any(probN > 1) or sum(probN) > 1 or probS > 1 or probS < 0 or abs(probS - sum(probN)) > 1e-3:
                        if warningSuppression == False:
                            print('Bad values at time t=' + str(
                                t[timePointIterator]) + ' lambda=' + str(
                                lambdaRange[lambdaIterator]) + ' rho=' + str(
                                rhoRange[rhoIterator]) + ' r=' + str(rRange[rIterator]))
                            print('any(probN>1=' + str(
                                any(probN > 1)) + ', sum(probN)>1=' + str(
                                sum(probN) > 1) + ', (probS-sum(probN))=' + str(
                                probS - sum(probN)))
                        nBadValues += 1
                    else:
                        if warningSuppression == False:
                            print('Completed time t=' + str(t[timePointIterator]) + ' weeks, lambda=' + str(lambdaRange[lambdaIterator]) + ' rho=' + str(rhoRange[rhoIterator]) + ' r=' + str(rRange[rIterator]))

                    PScanPP_local[rIterator][rhoIterator][timePointIterator][:] = probN / probS
                PScanPP_local[rIterator][rhoIterator] = calculateLikelihood(
                    PScanPP_local[rIterator][rhoIterator], data)
        PSurvScanPP[lambdaIterator] = PSurvScanPP_local
        PScanPP[lambdaIterator] = PScanPP_local


    return np.array(PScanPP[0])


def calculateAnalyticalIndividualTimepointsLikelihood(t, nRange, lambdaRange,
                                                      rhoRange, rRange, data):
    maxN = max(list(map(max, nRange)))
    lamda = lambdaRange[0]
    timePoints = range(len(t))
    PScanPP = [[np.zeros(len(rRange)) for L2 in range(len(rhoRange))] for L4 in
               timePoints]
    nBadValues = 0
    
    for timePointIterator in timePoints:
        for rhoIterator in range(len(rhoRange)):
            for rIterator in range(len(rRange)):
                probS = Psurv(t[timePointIterator], lamda, rRange[rIterator], rhoRange[rhoIterator])
                avgN = 1 / probS
                Niter = max([normal_round(int(4 * avgN)),
                             normal_round(1.5 * max(nRange[timePointIterator])), 10])
                probN = Pn(nRange[timePointIterator], t[timePointIterator], lamda, rRange[rIterator], rhoRange[rhoIterator],
                           Niter, 'returnConsecutive', maxN)

                if any(probN > 1) or sum(probN) > 1 or probS > 1 or probS < 0 or abs(probS - sum(probN)) > 1e-3:
                    if warningSuppression == False:
                        print('Bad values at time t=' + str(t[timePointIterator]) + ' rho=' + str(
                            rhoRange[rhoIterator]) + ' r=' + str(rRange[rIterator]))
                        print('any(probN>1=' + str(
                            any(probN > 1)) + ', sum(probN)>1=' + str(
                            sum(probN) > 1) + ', (probS-sum(probN))=' + str(
                            probS - sum(probN)))
                    nBadValues += 1
                else:
                    if warningSuppression == False:
                        print(
                            'Completed time t=' + str(t[timePointIterator]) + ' weeks, rho=' + str(
                                rhoRange[rhoIterator]) + ' r=' + str(rRange[rIterator]))

                sizeProb = probN / probS
                PScanPP[timePointIterator][rIterator][rhoIterator] = calculateLikelihood(sizeProb, data[timePointIterator])
    return PScanPP

# %% [markdown]
## Search for the most likely r, ρ parameter sets

Call necessary functions to trigger analysis based on the user's specified approach

# %% [code]
def getCloneSizeRange(counts):
    cloneSizeRange = []
    for cloneSizeDist in counts:
        cloneSizes = [size + 1 for size, freq in enumerate(cloneSizeDist) if freq != 0]
        cloneSizeRange.append(list(range(min(cloneSizes),max(cloneSizes))))
    return cloneSizeRange

logLikelihoods = []

if analysis_plan['method'] == 'simulation':
    nSize = len(np.transpose(counts))
    #Binning of clone sizes
    nBin = int(round(np.log(nSize)/np.log(2)+0.5)) 
    cloneSizeBins = np.append([1],[2**(n-1) for n in range(1,(nBin+1))]) 
    expRate = ExponentialGenerate(102, meanDivisionTimeInDays)
    logLikelihoods = calculateSimulationLikelihood(expRate, meanDivisionTimeInDays,
                                            counts, times, cloneSizeBins)
    
elif analysis_plan['method'] == 'analytical':
    t = times / 7
    meanDivisionRatePerWeek = 7 / meanDivisionTimeInDays
    lambdaRange = [meanDivisionRatePerWeek]
    if analysis_plan['individual_timepoints']:
        nRange = getCloneSizeRange(counts)
        logLikelihoods = calculateAnalyticalIndividualTimepointsLikelihood(t, nRange,
                                                                    lambdaRange,
                                                                    rhoRange,
                                                                    rRange,
                                                                    counts)
    else:
        nRange = [list(range(1, max(np.where(t > 0)[0]) + 1)) for t in counts]
        logLikelihoods = calculateAnalyticalLikelihood(t, nRange, lambdaRange,rhoRange, rRange, counts)

# %% [markdown]
### Plotting

Parameter inference results are plotted here. Results are visualised as heatmaps of *r* and *ρ* likelihoods.

# %% [code]
def plotLikelihood(result, title):
    plt.imshow((np.flip(np.transpose(result), axis=0)), cmap='coolwarm',
               interpolation='nearest', extent=[max_r/parameterBins, max_r, max_rho/parameterBins, max_rho],
               aspect=max_r/max_rho)
    plt.colorbar()
    plt.title(title)
    plt.xlabel('r')
    plt.ylabel(r'$\rho$')
    plt.show()
    
if not analysis_plan['individual_timepoints']:
    logLikelihoods = np.expand_dims(logLikelihoods,axis=0) # add extra dimension (fake timepoint) for plotting purposes 
    
if analysis_plan['method'] == 'simulation' and analysis_plan['individual_timepoints']:
    logLikelihoods = np.transpose(logLikelihoods, (2, 0, 1)) # swap [r][rho][t] numpy array to [t][r][rho] for plotting purposes


for timepoint in range(len(logLikelihoods)):
    if analysis_plan['individual_timepoints']:
        t = "t" + str(timepoint)
    else:
        t = 'all'

    plotLikelihood(logLikelihoods[timepoint], "Loglikelihood " + t)
    scaledR = logLikelihoods[timepoint] - max(max(r) for r in logLikelihoods[timepoint])
    expR = 10 ** (scaledR)
    plotLikelihood(expR, "Likelihood " + t)

# %% [markdown]
### Calculate 95% confidence intervals

parameter 95% confidence intervals are calculated and *r* and *ρ* likelihoods are plotted separately.

# %% [code]
def calculateConfidenceIntervals(parameterRange,parameterNormalised):
    parameterDistribution = np.cumsum(parameterNormalised)
    commonIndices = np.intersect1d(np.where(parameterDistribution > 0.025),np.where(parameterDistribution < 0.975))
    if len(commonIndices) > 0 :
        minIndex = np.amin(commonIndices)
        maxIndex = np.amax(commonIndices)
        ci = [parameterRange[minIndex], parameterRange[maxIndex] ]
    else:
        ci = 'Unable to estimate 95% CI'

    return ci


for timepoint in range(len(logLikelihoods)):
    if analysis_plan['individual_timepoints']:
        t = "t" + str(timepoint)
    else:
        t = 'all'
    scaledR = logLikelihoods[timepoint] - max(max(r) for r in logLikelihoods[timepoint])
    expR = 10 ** (scaledR)
    allSum = sum([sum(r) for r in expR])
    rNormalised = [sum(r) for r in expR]/allSum
    rhoNormalised = [sum(rho) for rho in np.transpose(expR)] / allSum
    print('timepoint: ' + t)
    print('r 95% CI: ' + str(calculateConfidenceIntervals(rRange,rNormalised)))
    print('rho 95% CI: ' + str(calculateConfidenceIntervals(rhoRange,rhoNormalised)))


plt.step(rRange, rNormalised)
plt.xlabel('r')
plt.ylabel('Likelihood')
plt.show()

plt.step(rhoRange, rhoNormalised)
plt.xlabel('rho')
plt.ylabel('Likelihood')
plt.show()

# %% [code]
```
