## Supplementary material for "Methods for analysing lineage tracing datasets": Jupyter notebooks for analysis: README.txt

We provide two notebooks that contain different workflows for parameterising lineage tracing clonal data. We present the SMC-ABC method as the most appropriate for accurate and efficient analysis of lineage tracing datasets.


SMC-ABC NOTEBOOK: 

In the SMC-ABC notebook we implement an Sequential Monte Carlo Approximate Bayesian Computation using a Non-Markovian simulator of the sinle progenitor model.

MLE NOTEBOOK:

The MLE notebook allows for applying Maximum Likelihood Estimation to find the most likely parameter combination using either a) Gillespie simulations of the single progenitor model or b) an analytical formula that computes parameter probabilities.

DEPENDENCIES:
In order to run the analysis workflows provided by this notebook, you need to have Python 3 and Jupyter Notebook installed. 

the following libraries need to be installed:

numpy 1.17.2
pandas
scipy
matplotlib
pyabc (for the SMC-ABC notebook only)
mpmath (for the MLE notebook only)


USAGE:

Please provide your input data file and your preferred analysis parameters at the "User configuration" block of each notebook. A brief explanation on each parameter is provided there. We provide an example input data file to use it as a template. In order to run the workflows successfully all code blocks must be executed sequentially. 
