## Supplementary figures and images for "Methods for analysing lineage tracing datasets"

### Cartoon.png

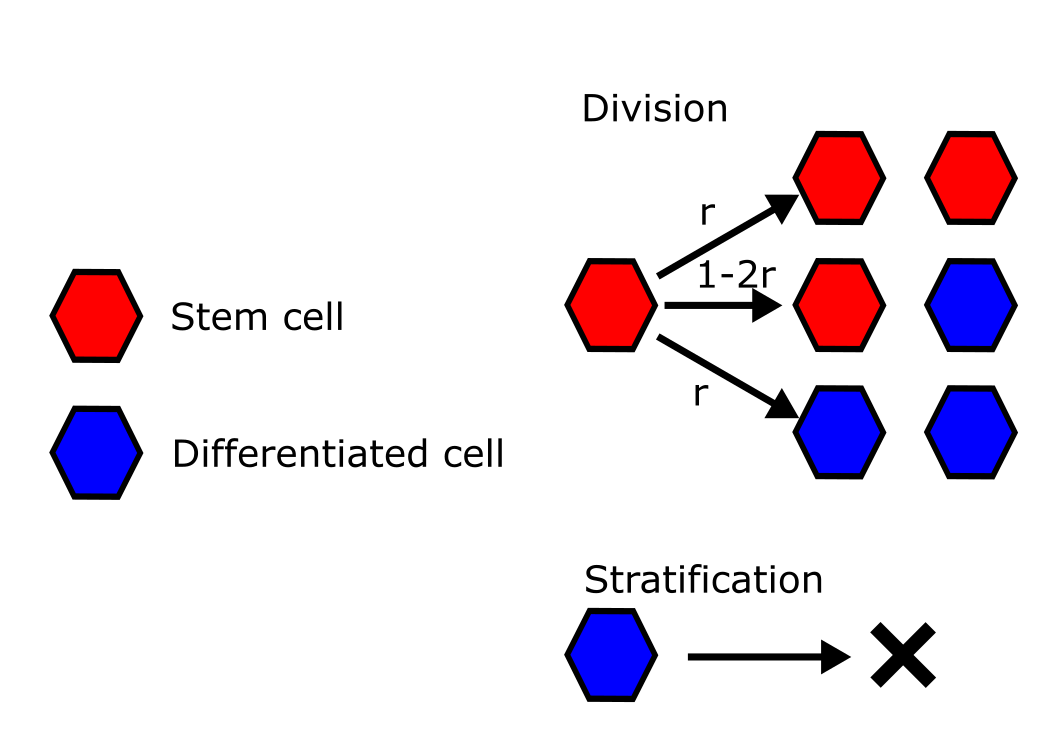

### CellFate.png

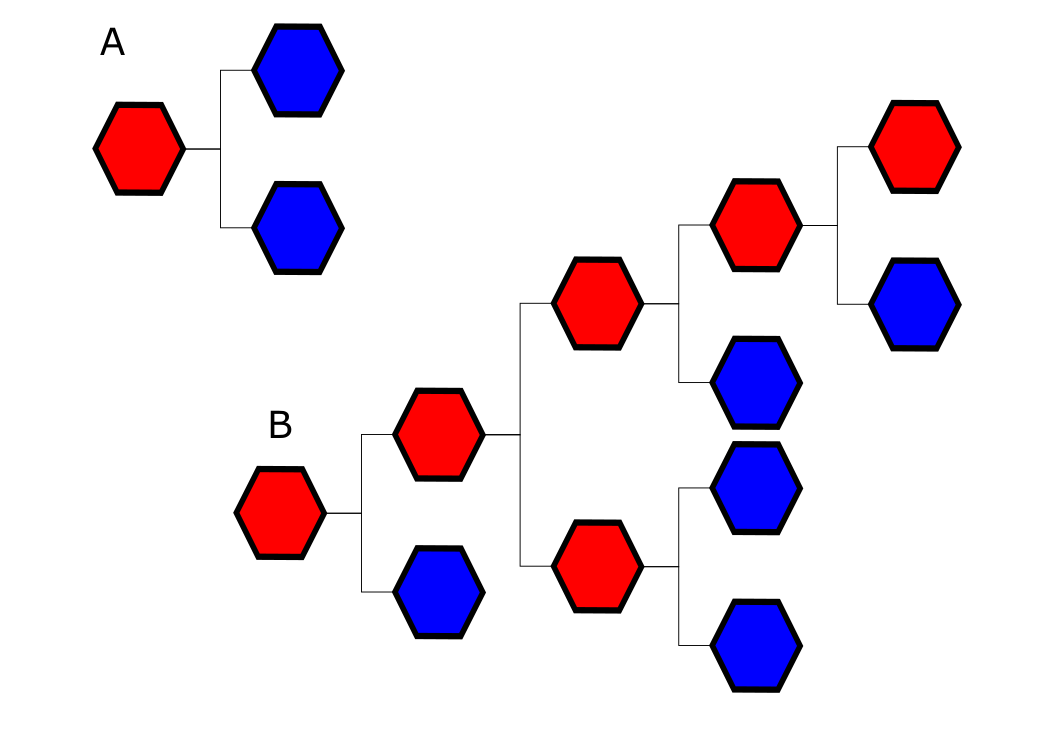
